# Health care seeking behavior and associated factor among mothers/caregivers of under-five children with acute diarrhea in Dangila zuria Woreda, North west Ethiopia

**DOI:** 10.1101/667923

**Authors:** Biresaw Nega, Kassawmar Angaw Bogale, Zelalem Mehari Nigussie

**Affiliations:** Master of public health, Awi zonal health department, Ethiopia; Master of pubic health, Department of Epidemiology and Biostatistics, School of Public Health, College of Medicine and Health Sciences, Bahir Dar University, Bahir Dar, Ethiopia; Master of Sciences in Biostatistics, Department of Epidemiology and Biostatistics, School of Public Health, College of Medicine and Health Sciences, Bahir Dar University, Bahir Dar, Ethiopia

**Keywords:** Health care Seeking Behavior, Under five children’s, diarrhea, mother’s caregiver

## Abstract

**Introduction:** Health care seeking interventions can reduce child mortality from easily treatable diseases, however, significant numbers of children die by diarrheal disease due to delays in seeking care in Ethiopia. Thus, the aim of this study was assessed health care seeking behavior and associated factors among mothers/caregivers of under-five children with acute diarrhea in Dangila zuria Woreda, North West Ethiopia, 2018.

**Method:** Community-based cross-sectional study design was conducted using structured questionnaires. Data were entered Epi Data Version 3.1 and analyzed using SPSS Version 23. Descriptive statistics were used to examine patterns of health care seeking behavior and multivariable logistic regression analysis was applied to identify factors associated with health care seeking behavior

**Result:** The magnitude of health care seeking behavior of mothers/caregivers of under-five children with acute diarrhea was found to be 77.7%. Primary level of education [AOR = 2.0; 95% (CI (1.1-3.9)], sex of child (male) [AOR = 1.7; 95% (CI 1.1-2.6)], availability of health facilities within 60 minutes walking distance [AOR = 2.4; 95 %(CI 1.4-4.1)], severity of illness [AOR=7.5; 95% (CI 3.7-15.2)], two or more under-five children in the family [AOR=0.6;95% (CI 0.4-0.9)], perceived cause of acute diarrhea, (new teeth [AOR =0.3;95% (CI 0.2-0.5)] were significantly associated with health care seeking behavior of mothers/caregivers.

**Conclusion:** Increasing the proximity of health facilities and educate mothers/caregivers about the importance of health care seeking behavior and cause of acute diarrhea were recommended to improve health care seeking behavior.

## Introduction

Health-seeking behavior has been defined as a sequence of remedial actions that individuals undertake to rectify perceived ill-health. It is the process of care seeking by individuals for improving the perceived disease and it is a reflection of the prevailing conditions, which interact synergistically to produce a pattern of care seeking but which remains fluid and therefore amenable to change(1).

Although under-five mortality has been reduced globally, diarrhea is the first leading cause of under-five children mortality in the world (2) and the situation in sub-Saharan Africa is still a major concern(3). Sub-Saharan Africa (SSA) has made the least progress in terms of reduction of child mortality compared to all other regions in the world(4). Health care seeking interventions have the potential to reduce child mortality, however, in developing countries, a large number of children die without ever reaching a health facility and (5).

World Health Organization reported that 70% of child deaths are related to delayed care-seeking(6). A study done in Nairobi, Kenya showed that healthcare-seeking practices for diarrhea remain great challenges and more than half (55%) of the caregivers seeking inappropriate health care with a large number of caregivers (35%) taking no action regarding the child diarrheal(7). Similarly, a study done in Tanzania indicated that only 23.0 % of children with acute diarrhea treated at a health facility(8).

Poor health care seeking behavior increases the chance of morbidity and mortality of children (9). Significant numbers of children die due to delays in seeking care in Ethiopia (10). For instance, a study done in Bahir Dar city indicated that diarrhea was higher in under-five children but treatment seeking was low due to low seeking behavior of mothers(11).

In the study area, the woreda Health Management Information System (HMIS) report indicated that, among expected diarrheal cases, only 27% of under-five children were treated in 2017 (12). However, factors to hinder health care seeking were unknown.

Even if, little studies were conducted in Ethiopia regarded health care seeking behaviors for acute diarrhea, previous studies were not community-based and disease-specific. Therefore, the aim of this study was to assess health care seeking behavior and associated factors among mothers/caregivers of under-five children with acute diarrhea in Dangila Zuria Woreda, North West Ethiopia, 2018. This study might help program planners and implementers to design intervention pertaining to health care seeking behavior promotion, to treat diarrhea disease of under-five children and to this study may use as baseline for researchers.

There are two major frameworks that have been proposed to explain the health care services utilization of an individual from the behavioral aspect. These are Andersen & Newman model and the Kroeger’s model(13). Andersen’s health behavior model is common and used to study the determinants of health care seeking behavior. We used this model as a conceptual framework for our analyses(14).

According to Andersen’s behavioral model, predisposing, enabling, and need factors at the individual and community levels are instrumental for increasing health-seeking behavior and health facilities utilization. Predisposing factors were reflecting the families which are likely to use health services while enabling factors are those which promote or hinder health service use. The first two sets of factors are not enough until the family perceives the severity of the illness. This is called as ‘need’ factor which is the most immediate reason for health service use(15, 16). Inappropriate health care seeking behavior increases the chance of morbidity and mortality of children. So to improve the health of under-five age children, the health care seeking behavior of mothers/caregivers plays a major role in determining the infant’s survival(9).

**Fig. 1.**
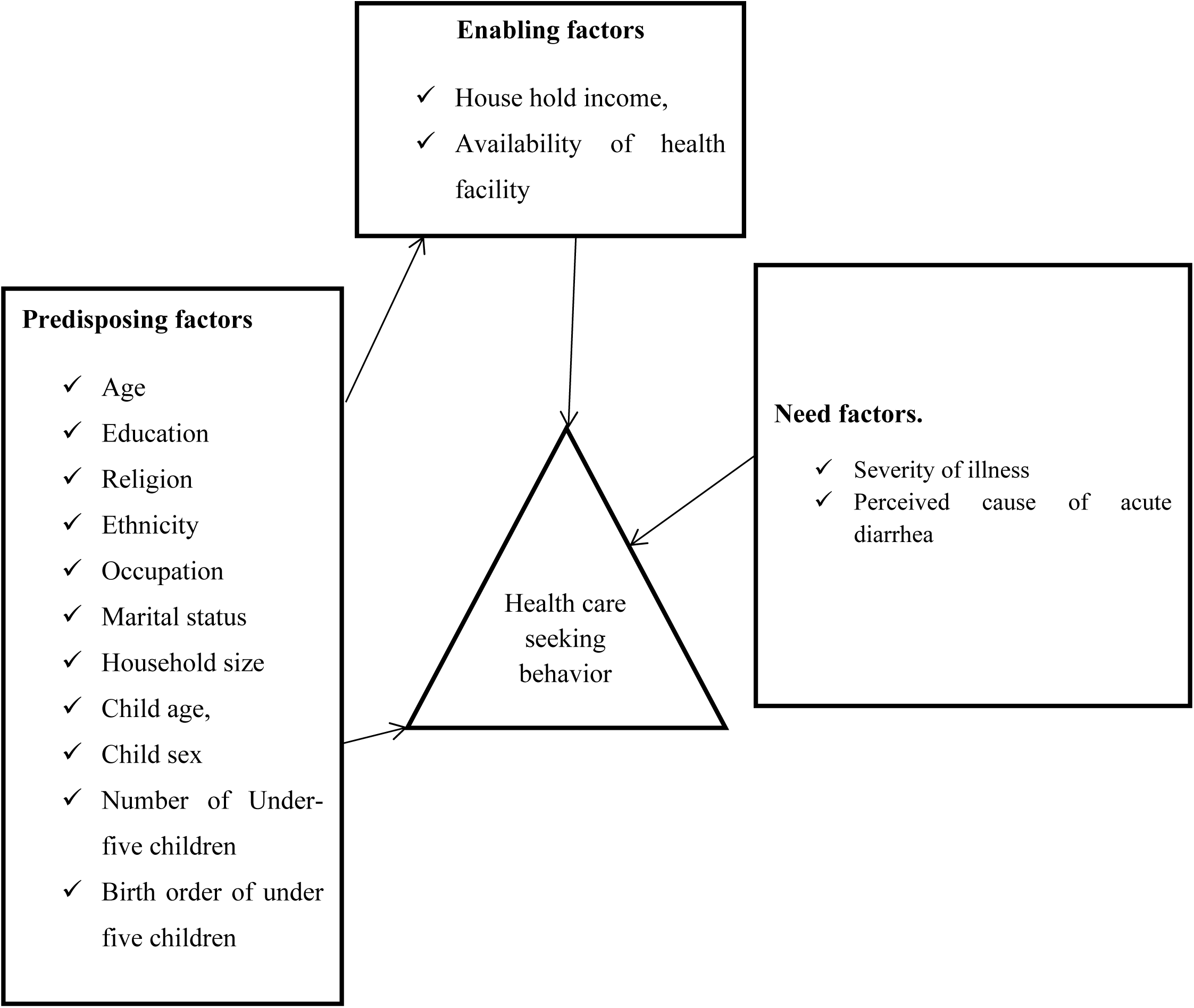
Conceptual framework for health care seeking behavior from literature review and modified from Andersen’s behavioral mode (19).

## METHODS

### Study settings

The study was conducted in Dangila zuria Woreda, Awi zone, Amhara Regional State, Ethiopia. Dangila Woreda is located 485 kilometers (k.ms) Northwest from Addis Ababa and 78 km from Bahir Dar, the capital city of the Amhara region. The Woreda has a total of 2 urban and 27 rural kebeles. According to information obtained from the Woreda office, it has a total of 36,749 households and 158, 022 populations out of these 21,396 were under-five children. The Woreda has six health centers, 27 health post and five private clinics(17).

### Study Period and Population

The study was conducted from November 1,2018 to December 1,2018. The source populations were all mothers/caregivers who had under-five children with acute diarrhea in the last 2 weeks of the preceding data collection period in Dangila Zuria Woreda.

Study Population Randomly selected mothers/caregivers from 10 kebeles who had under five children with acute diarrhea in the last 2 weeks of data collection period in Dangila Zuria Woreda.

### Inclusion and Exclusion Criteria

Inclusion Criteria all mothers/caregivers who had under-five children (1-59 months) with acute diarrhea in the last 2 weeks of data collection period and resided for at least 6 months in the study area.

Exclusion Criteria Mothers/caregivers who can’t communicate due to severe illness or sickness.

### Sample Size Determination

For sample size estimation, we assumed a 95% confidence interval (CI), a margin of error of 5%, a proportion of 56 % for health care seeking conducted nationally in four region of rural Ethiopia (Amhara, SNNP, Oromia and Tigray)(18), and a design effect of 1.5. As the source population was less than 10,000; sample size correction was performed. Then, 10% non-response rate was added to obtain the enough sample size

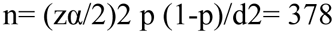

**n** = required sample sizes

When we consider the design effect (**1.5**), the sample size required became:

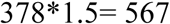

For possible nonresponse during the survey the final sample size is increased by 10% to n = 567 +10% which is 57 = 624. Therefore, the total sample size including 10% NRR was 624.

### Dependent variables

Health care seeking behavior

### Independent Variable

Predisposing factors: (socio-demographic factors: -mother/caregiver age, education, religion, ethnicity, occupation, marital status, household size, child age, child sex, no. of <5 children, birth order of under-five children. Enabling factors: (Household income, availability of governmental or private health facility

Need factors :(Severity of illness, Perceived cause of acute diarrhea)

### Sampling procedure

A total of 10 kebeles were included in the study, one kebeles in urban and 9 kebeles in rural were selected using the lottery method. Then systematic random sampling was employed to selected households after allocating the number of under-five children with acute diarrhea in each kebele proportionally. The first household was chosen near the Kebele Council as a starting point. Households with no under five children with acute diarrhea were excluded from the study. After a successful interviewee of each household the interviewer continue to the right side of the next household whenever there is more than one under five children with acute diarrhea in a household only one was selected using lottery method and all caregivers in the selected kebele that full fill the sampling criteria are interviewed until the required sample size was reached.

### Operational definition

#### Modern health care seeking behavior

Is care sought from health facilities; hospitals, health centers, private clinics or health posts for under-five children with acute diarrhea within 2 weeks (10).

#### Acute diarrhea

The passage of three or more abnormally loose watery or liquid stools over 24 h periods within 2 weeks prior to the data collection.

#### The severity of illness

Was identified simply according to mothers /caregiver’s perception of the disease condition.

### Data collection procedures and data quality control

Data were collected using questionnaires and carried out by face to face interview method. Data collectors and supervisor were coming from other non-selected and trained for two days on objectives, interview method, confidentiality, and data quality control. Two BSC nurses as the data collectors and one health office as the supervisor have participated. Firstly, the questionnaire was designed in English and translated into local Amharic language by language expert then translated into English for the consistency of the questionnaire. The pre-test was conducted on 5% of the sample size. After pre-testing further adjustments to the data collection tool was made to improve the clarity, understandability, and simplicity of the messages. Check the completeness of questionnaires page by page before and after the data collection were conducted. When the data were collecting supervise consecutively each data collectors and prepare regular meetings each day. The interviewed questionnaires were reviewed, collected and checked by the supervisor and principal investigator daily.

### Data processing and analysis

Data were entered Epi Data Version 3.1 and analyzed using SPSS Version 23. The data were checked for completeness and cleaned for the errors during data entry. Descriptive statistics were used to examine patterns of health care seeking behavior and multivariable logistic regression analysis was applied to identify factors associated with health care seeking behavior. Finally, 95% confidence interval and a p-value less than 0.05 was considered as significant associated factors of health care seeking behavior.

### Ethical considerations

Ethical clearance was obtained from Institutional Review Board committee of Bahir Dar University college of Medicine and health sciences.

Then ethical approval letter was taken to the respective health facility managers and kebele administrators and written consent was obtained from participants after informing of the confidentiality of the information they give and will not be given to anyone without their consent. The name of participant did not document in the questionnaire. The participants will have as full right to withdraw from study at any time during interview.

## Results

### Socio demographic characteristics of respondents and children

In this study, 624 mothers/caregivers who had under-five years’ old children with acute diarrhea were interviewed with a response rate of 100%. Six hundred five (97%) were mothers of the selected child, 229 (37%) in the age group of above 35 years with a mean age of 32 and 554 (88.8%) were married. Regarding sex and age of the child, 51% of children were males and 35% of children were in the age group of 12 to 24 months. (Table 1).

**Table 1:**
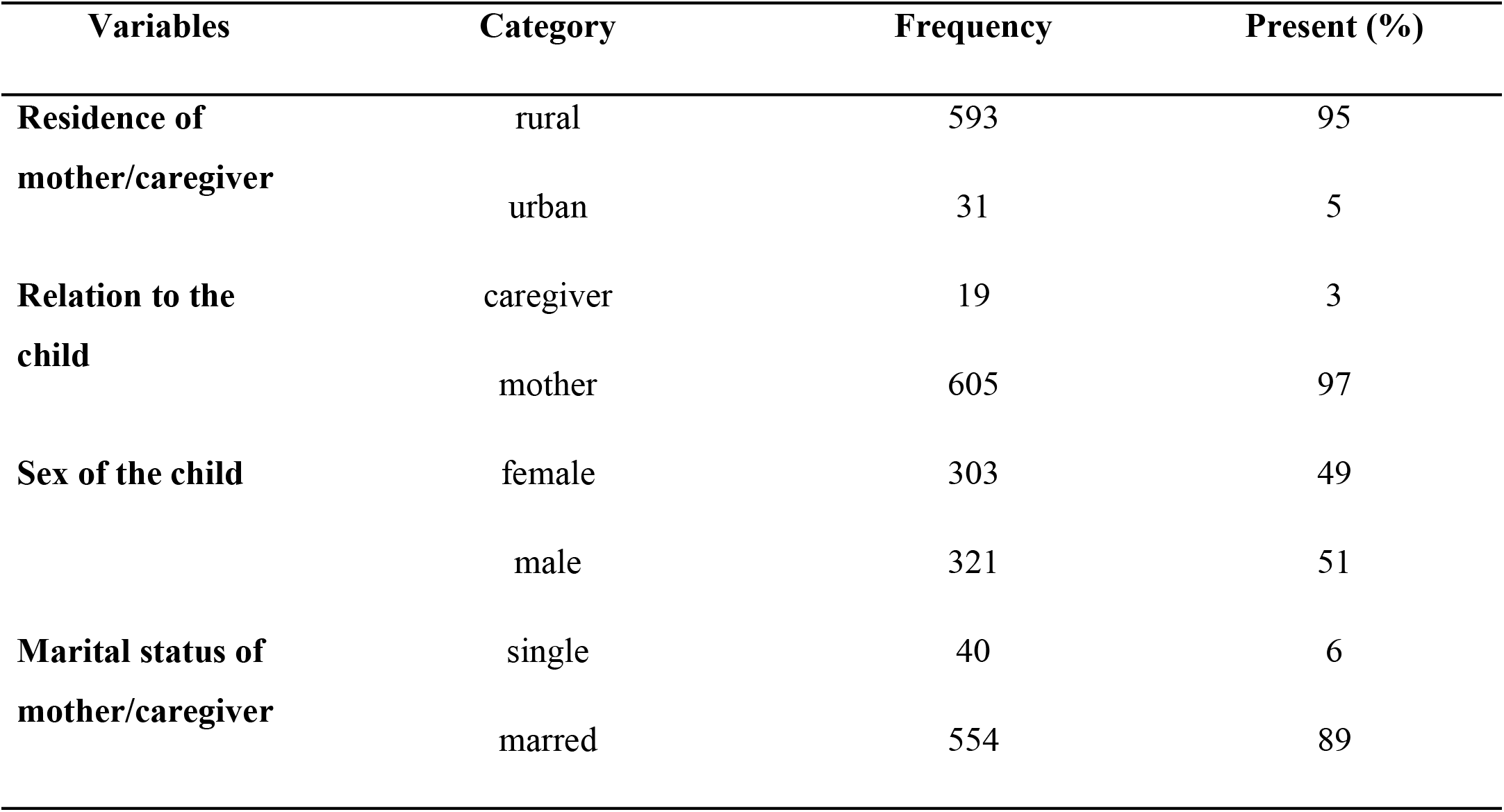
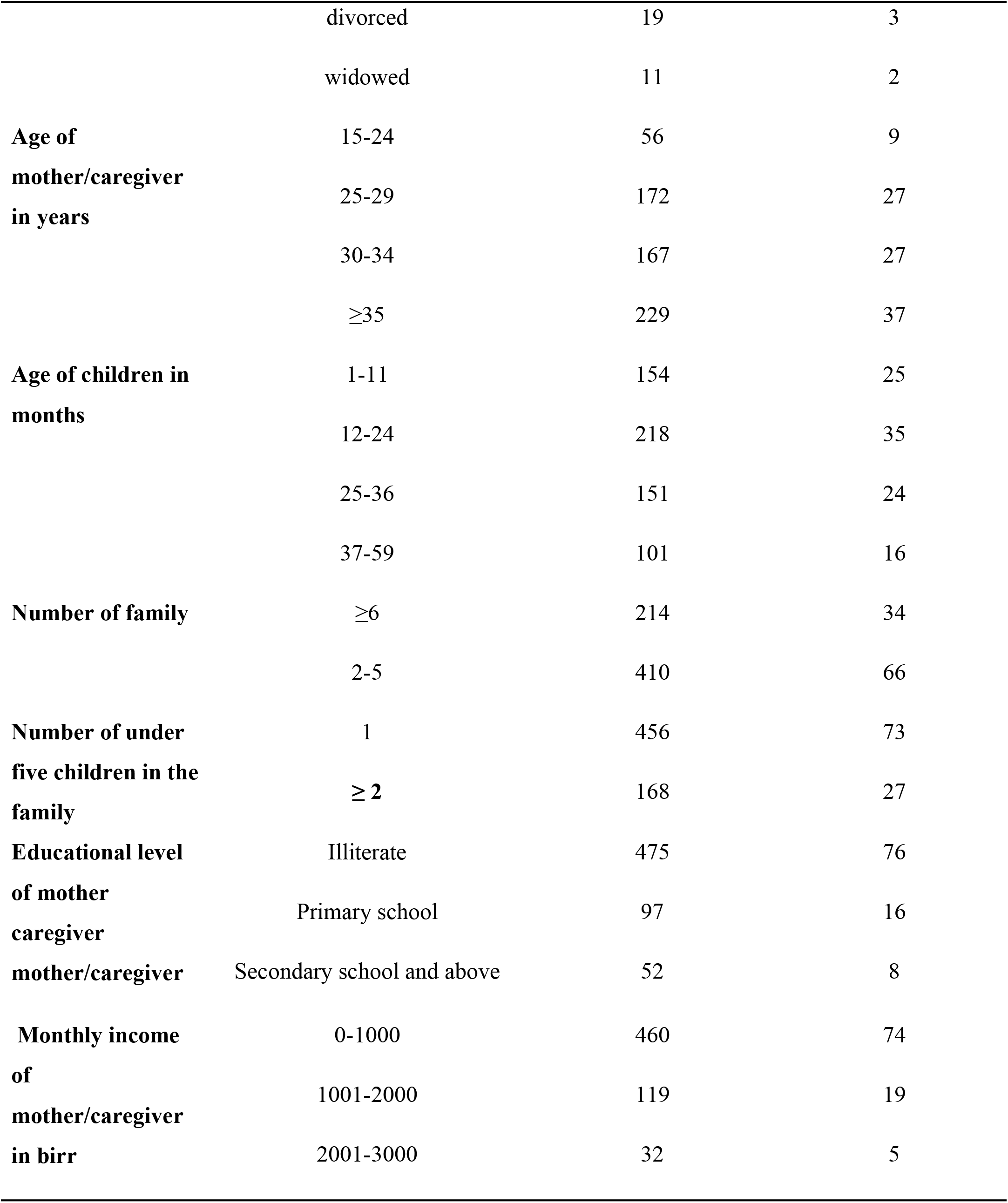
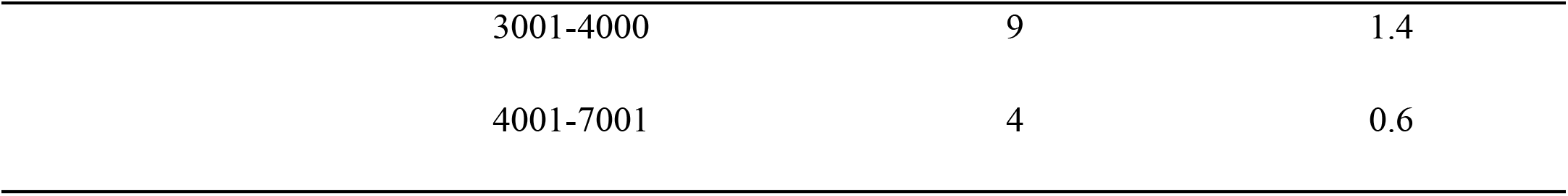
Socio demographic Characteristics of mothers/caregivers of under-five children with acute diarrhea in Dangila Woreda, 2018.

### Health care seeking behavior

Four hundred eighty-five (77.7%) (95%CI:74.4%-81.1%) mothers/caregivers of under-five children with acute diarrhea were sought health care from the health facility. Four hundred thirty-nine (90.5%) mothers/caregivers of under-five children with acute diarrheal were seen at government health facilities. The mean walking time to reach the health facilities for seeking care was 37.7(±21) min, and the mean number of elapsed days to get care after recognizing the sign of diarrhea was 2.5 (±3.4) days

### Reasons for not seeking care at the health facility

Among the reasons for not seeking health facilities, 72% of mothers/caregivers were not seeking care for their child due to illness was not sever and 36% were due to mothers/caregivers were busy. Among privet health facility utilizer, 60% of mothers/caregivers prefer private health facility due to treatment obtained within a short period of time (Table 2).

**Table 2:**
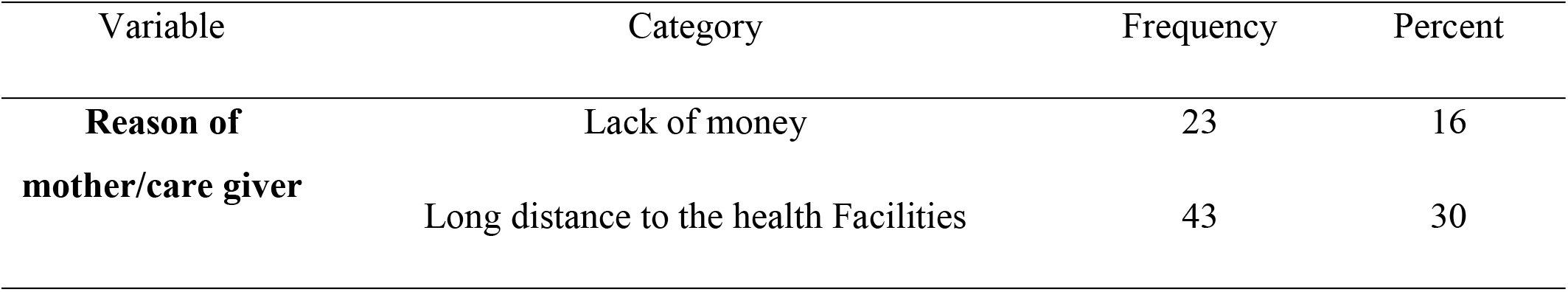
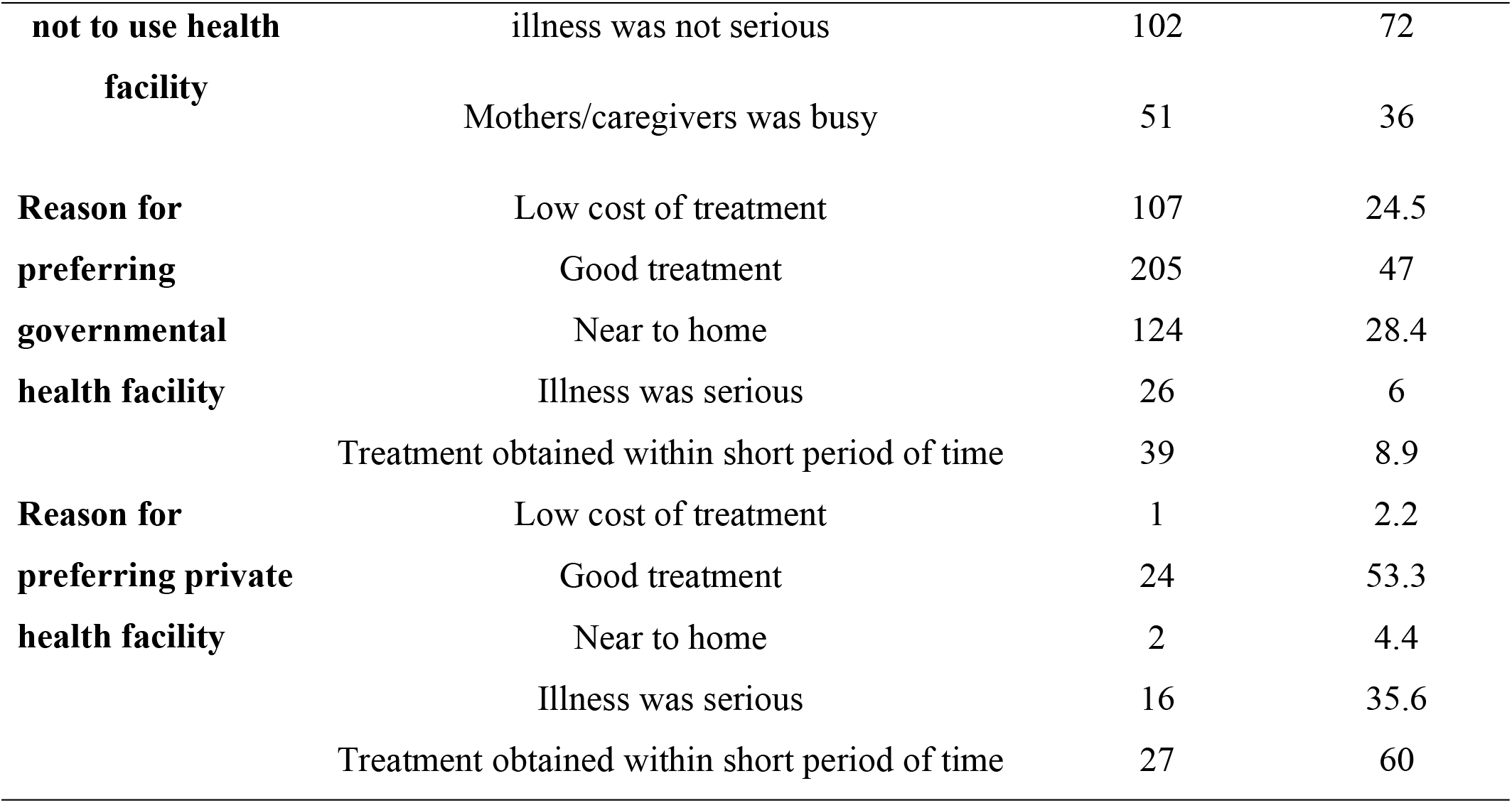
Health facilities utilization and reasons for not seeking care at health of mother/caregiver of under five children with acute diarrhea Dangila Woreda, 2018.

### Factors affecting health care seeking behavior

In bivariable logistic regression of predisposing factors like: mothers/ caregivers educational level of primary school, sex of child (male) and number of under five years old children in the family (2 or more), from enabling factors: availability of health facilities within 60 min walking distance and from the need factors: severity of disease and cause of acute diarrhea (those who don’t know the cause of acute diarrhea, poor hygiene and new teeth) were entered to multivariable logistic regression analysis.

In multivariable logistic regression analysis there was statistically significant association between health care seeking behavior of mothers/caregivers of under-five children with acute diarrhea and sex of child (male), educational level of mothers/caregivers who have primary level of education (1-8 grade), number of under-five children in the family (2 or more), availability of health facilities within 60 min walking distance and severity of illness according to their perception and perceived cause of acute diarrhea due to new teething. Those mothers/caregivers who had a male child were more likely to have health care seeking behavior of under-five children with acute diarrhea compared to those who had a female child. (AOR = 1.7; 95% CI (1.1– 2.6))

Those mothers/caregivers who had 2 or more under-five children in the family were 40% less likely to have health care seeking behavior compared to those who had one under five child in the family with acute diarrhea (AOR=0.6;95% CI (0.4-0.9)).

Those mothers/caregivers who had the educational levels of primary school were two times higher than those who were illiterate to have health care seeking behavior of under-five children with acute diarrhea (AOR = 2.0;95% CI (1.1–3.9)).

Those mothers/caregivers who had severely ill children according to their perception were nearly seven and half times higher than those who were not severely ill to have health care seeking behavior for under five-children with acute diarrhea (AOR=7.4; 95% CI (3.7-15.2))

Those mothers/caregivers who perceived new teething as the cause of acute diarrhea were 70 % less likely to have health care seeking behavior of under-five children with acute diarrhea as compared to those who did not perceive new teething as the cause of acute diarrhea (table 3).

**Table 3:**
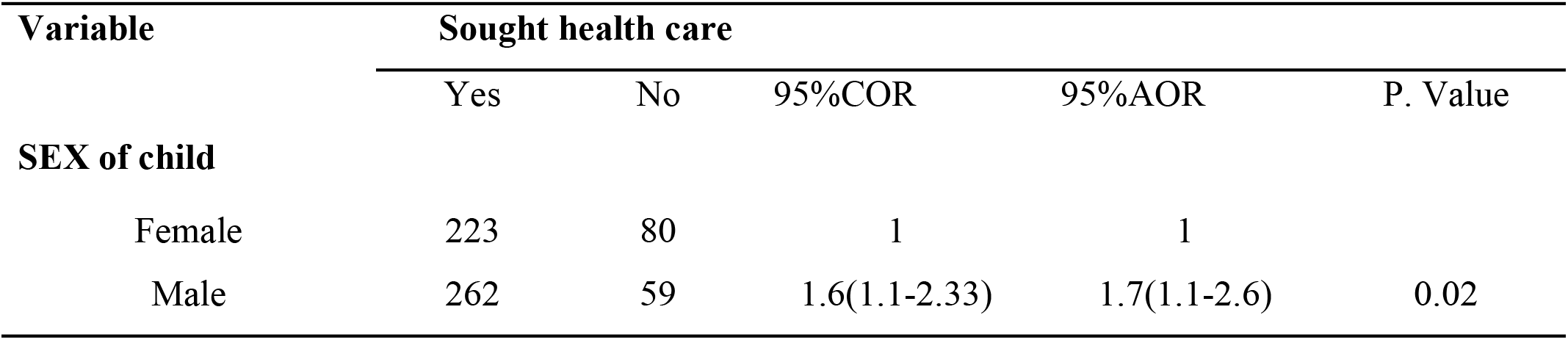
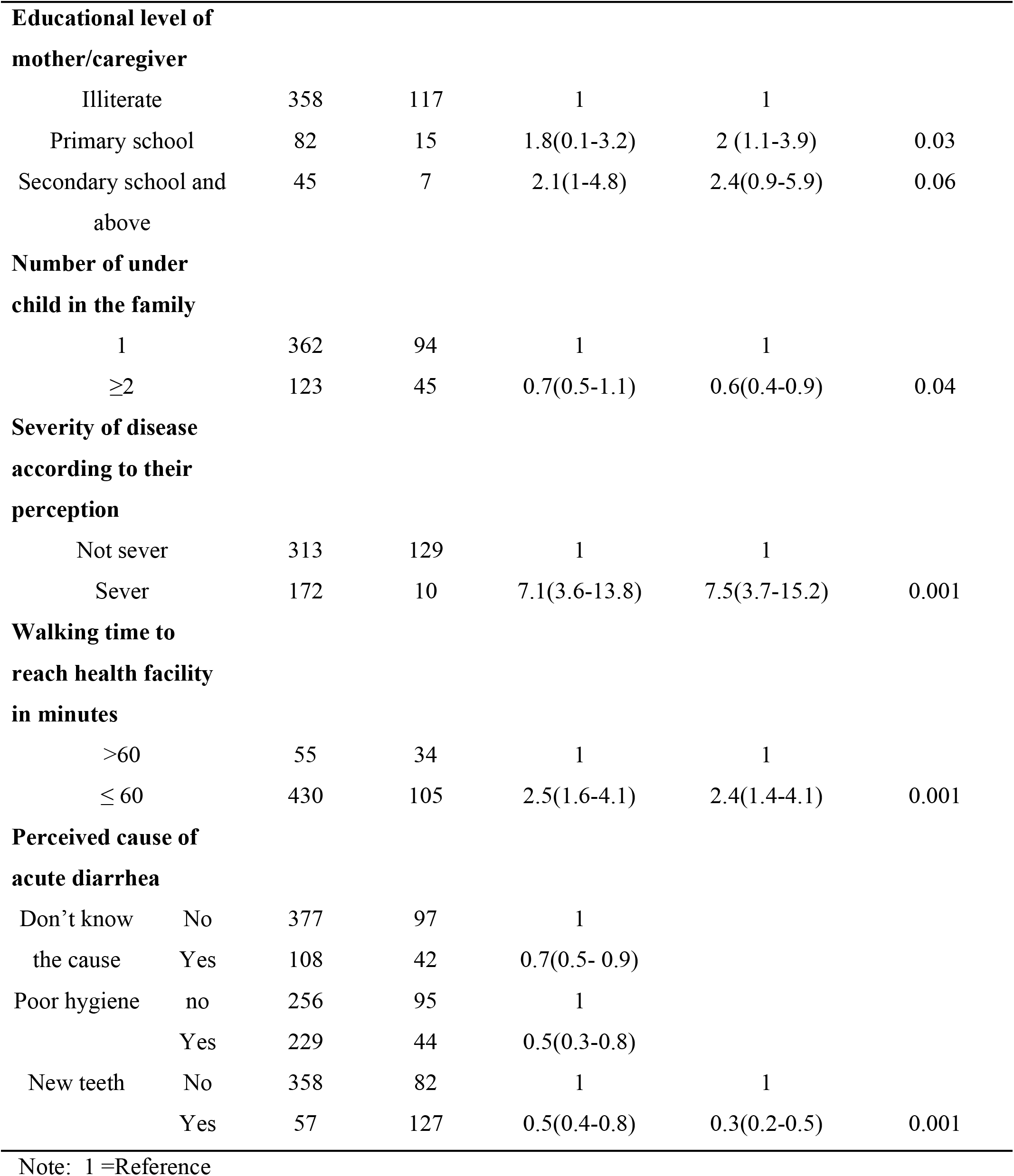
Factors associated with health-seeking behavior mother/caregiver of under five children with acute diarrhea Dangila Woreda, 2018.

## Discussion

The magnitude of health care seeking behavior of mothers/caregivers of under-five children with acute diarrhea was 77.7 % (95%CI: 74.4% - 81.1%). This is consistent with the study conducted in the Oromia region Jeldu district (81.1%) (10), and lower than the study conducted in the Gambia of Ethiopia (81.5%) (19) this might be due to the difference in geographic and socioeconomic conditions.

The magnitude of health care seeking behavior of this study was higher than the study conducted at the national level of Ethiopia (56%) (18), and in Addis Abeba (56.6%) (20). This difference might be due to seasonal variation.

In terms of sex preference in healthcare-seeking behavior, the study shows a significant difference between male and female children, a male child is 1.7 more likely to seek health care than female children. A number of studies has shown boys preference in seeking health care especially for childhood illnesses like acute diarrhea (21-23). The likely explanation for the difference in seeking treatment differently for boy and girl children could be due to cultural effect and gender inequality that cause disadvantages females in the community, which in turn, may impose mothers/caregivers to give priority to a boy child.

Those mothers who have more than or equal to two number of under-five children in the family members were 40% less likely to seek medical care than those who have one under-five child in family members, contrary to our finding, study done in Addis Ababa indicated that there was no difference in health care seeking behavior of mothers/caregivers of under-five children with acute diarrhea due to number of under-five children in the family (20). This might be due to the incapability of the mothers/caregivers to care for many under-five children and a shortage of money for seeking care in the study area as compared to Addis Ababa mothers/caregivers.

In this study, the relationship between the educational level of the respondents (primary level of education) and health care seeking behavior was found to be statistically significant. Which is consistent with studies done in Nigerian, Nepal, and Ensaro district North Shoa Zone (24-26). This is because educated mothers/caregivers can understand health education messages presented in mass media and through other methods more than the none educated ones.

Another finding from this research is that proximity to the health facility within 60 minutes walking time has significantly associated with the health-seeking behavior of the mother caregiver. This is similar to other studies done in the Oromia region (22, 27). This is because living far away from the location of health facility may force mothers to walk or travel long distance which could lead to tiredness, exposure to hazard or incurring more expenses. Due to this, the mothers might be discouraged from presenting their children for medical treatment.

In the current study, mothers who perceived illness was sever were seven times more likely to seek medical care than those who perceive the illness is not severed. This finding is similar to a study done in Mekele city (28). This may be due to the mother’s belief of illness will improve by itself and wait until their child shows another symptom. So, this could be an indicator of as mothers are being heath seeker while their child is seriously ill.

We found that the perceived cause of acute diarrhea (new teething) decreases the likelihood of seeking care. Contrary to this finding, some studies reported that the perceived cause of acute diarrhea was not a reliable predictor of care seeking by mothers/caregivers((20, 29). It might be due to cultural and socio-demographic differences.

### Limitations

The study utilized perception and oral report of an individual for health care seeking report which may be increase the magnitude of health seeking behavior. Qualitative study was not done to triangulate it.

### Conclusion and recommendations

Even if many mothers/caregivers of under-five children with acute diarrhea sought health care from a health facility, still there are the number of mothers/care givers of under-five children with acute diarrhea who did not seek medical care. Education of mother/caregiver, sex of the child, availability of health facilities within 60-minute walking distance, the severity of illness, number of under-five children in the family and perceived cause of diarrhea were the independent factors for health care seeking behavior of mothers /caregivers of under-five children with acute diarrhea.

Health care facilities should be brought close to inhabitants of remote areas through establishment of health facility. Health workers should educate the mother/caregiver about the importance of health care seeking behavior especially about the severity of acute diarrhea and the cause of acute diarrhea. Similarly, the health bureau with the collaboration of the education bureau and other sectors should promote female education and empowerment to balance gender inequality. Further research that can minimize this limitation should be done to understand the health care seeking behavior of mothers/caregivers of under-five children with acute diarrhea.

## ACKNOWLEDGEMENT

First and for most, we would like to forward the deepest appreciation and thanks to Dangila Zuria Woreda health office and Health Extension Workers for their valuable information provided to us. Our special appreciation go to study participants, data collectors, and supervisors, and also we would like to Bahir Dar University college of Medicine and health Sciences, Department of Epidemiology and Biostatistics for giving this chance.

## REFERENCES

1. Atashbahar O, Bahrami MA, Asqari R, Fallahzadeh H. An examination of treatment seeking behavior affecting factors: a qualitative study in Iran. World Applied Sciences Journal. 2013;25(5):774–81.

2. UNICEF&WHO. Diarrhea: Why children are still dying and what can be done. The Lancet. 2012.

3. UNICEF. Levels and Trends in Child Mortality. 2015.

4. Chopra M, Mason E, Borrazzo J, Campbell H, Rudan I, Liu L, et al. Ending of preventable deaths from pneumonia and diarrhoea: an achievable goal. The Lancet. 2013;381(9876):1499–506.

5. De Souza AT, Peterson K, Andrade F, Gardner J, Ascherio A. Circumstances of post-neonatal deaths in Ceara, Northeast Brazil: mothers’ health care-seeking behaviors during their infants’ fatal illness. Social science & medicine. 2000;51(11):1675–93.

6. Kigali RCU, Angelique Charlie Karambizi, Lisine Tuyisenge, Peter Cartledge. The Pan African Medica Journa. 2018.

7. Illness Health care-seeking practices of caregivers of under-five children with diarrheal diseases in two informal settlements in Nairobi, Kenya.

8. Kanté AM, Gutierrez HR, Larsen AM, Jackson EF, Helleringer S, Exavery A, et al. Childhood illness prevalence and health seeking behavior patterns in rural Tanzania. BMC public health. 2015;15(1):951.

9. D’souza RM. Care-seeking behavior. Clinical infectious diseases. 1999;28(2):234-.

10. Kolola T, Gezahegn T, Addisie M. Health care seeking behavior for common childhood illnesses in Jeldu District, Oromia Regional State, Ethiopia. PloS one. 2016;11(10):e0164534.

11. Awoke W. Prevalence of childhood illness and mothers’/caregivers’ care seeking behavior in Bahir Dar, Ethiopia: A descriptive community based cross sectional study. Open Journal of Preventive Medicine. 2013;3(02):155.

12. Dangila Woreda. Health Mangment Information System(HMIS) report of under-five children disease report. dangila woreda; 2017.

13. K.B DC. Health care seeking behaviour. A theoretical perspective Pushpalata N kanbarkar Department of sociology, Rani channamma University, Belgavi Karnataka Dr Chandrika KB Department of sociology, Rani channamma University, Belgavi Karnataka. 2015.

14. Bahrami MA, Atashbahar O, Shokohifar M, Montazeralfaraj R. Developing a valid tool of treatment seeking behavior survey for Iran. J Novel Appl Sci. 2014;3(6):651–60.

15. Andersen R. A behavioral model of families’ use of health services. A behavioral model of families’ use of health services. 1968(25).

16. Andersen R NJ. Societal and individual determinants of medical care utilization in the United States. Milbank Mem Fund Q Health:. 1973;51(1):95–124.

17. Dangila Woreda. performance report and health profile 2018.

18. Addis Ashenafi AMK, Ameha A, Erbo A, Getachew N, Betemariam W. Effect of the health extension program and other accessibility factors on care-seeking behaviors for common childhood illnesses in rural Ethiopia. Integrated Community Case Management (iCCM) at Scale in Ethiopia: Evidence and Experience. 2014;52:57.

19. Saha D, Akinsola A, Sharples K, Adeyemi MO, Antonio M, Imran S, et al. Health Care Utilization and Attitudes Survey: understanding diarrheal disease in rural Gambia. The American journal of tropical medicine and hygiene. 2013;89(1_Suppl):13–20.

20. Adane M, Mengistie B, Mulat W, Kloos H, Medhin G. Utilization of health facilities and predictors of health-seeking behavior for under-five children with acute diarrhea in slums of Addis Ababa, Ethiopia: A community-based cross-sectional study. Journal of Health, Population and Nutrition. 2017;36(1):9.

21. Sudharrsanam M, & Rotti, S. Factors determinig helath seeking behaviour for sick children in a fisherman community in Podicherry. Indian Journal on Community Medicine. 2007.

22. Degefa G, Gebreslassie M, Meles KG, Jackson R. Determinants of delay in timely treatment seeking for diarrheal diseases among mothers with under-five children in central Ethiopia: A case control study. PloS one. 2018;13(3):e0193035.

23. Khera R, Jain S, Lodha R, Ramakrishnan S. Gender bias in child care and child health: global patterns. Archives of disease in childhood. 2013:archdischild-2013-303889.

24. Sreeramareddy CT, Shankar RP, Sreekumaran BV, Subba SH, Joshi HS, Ramachandran U. Care seeking behaviour for childhood illness-a questionnaire survey in western Nepal. BMC international health and human rights. 2006;6(1):7.

25. Abdulraheem I, Parakoyi D. Factors affecting mothers’ healthcare-seeking behaviour for childhood illnesses in a rural Nigerian setting. Early Child Development and Care. 2009;179(5):671–83.

26. Shewasinad S. Assessment of Mothers/Care Givers Health Care Seeking Behavior for Childhood Illness in Rural Ensaro District, North Shoa Zone, Amhara Region, Ethiopia 2014: Addis Ababa University; 2014.

27. Assefa T, Belachew T, Tegegn A, Deribew A. MOTHERS’HEALTH CARE SEEKING BEHAVIOR FOR CHILDHOOD ILLNESSES IN DERRA DISTRICT, NORTHSHOA ZONE, OROMIA REGIONAL STATE, ETHIOPIA. Ethiopian Journal of Health Sciences. 2008;18(3).

28. Fissehaye T, Damte A, Fantahun A, Gebrekirstos K. Health care seeking behaviour of mothers towards diarrheal disease of children less than 5 years in Mekelle city, North Ethiopia. BMC research notes. 2018;11(1):749.

29. Hustache S. Health care seeeking behavior for diarrhea in under 5 children in rural Niger. biomed central 2011.

